# Chemical biofilm dislodgement with chelating and reducing agents in comparison to sonication: implications for the diagnosis of implant associated infection

**DOI:** 10.1101/604637

**Authors:** Svetlana Karbysheva, Maria Eugenia Butini, Mariagrazia Di Luca, Tobias Winkler, Michael Schütz, Andrej Trampuz

## Abstract

Sonication of removed devices improved the microbiological diagnosis of infection. Recently, chemical agents have been investigated for dislodgement of biofilms, including the chelating agent ethylenediaminetetraacetic acid (EDTA) and the reducing agent dithiothreitol (DTT). We compared the efficacy of chemical methods (EDTA and DTT) to sonication for biofilm dislodgement. *Staphylococcus epidermidis* (ATCC 35984) and *Pseudomonas aeruginosa* (ATCC 53278) biofilms were grown on porous glass beads for 3 days. After biofilm formation, beads were exposed to 0.9% saline, sonication and/or chemical agents. Quantitative and qualitative biofilm analyses were performed by colony counting (CFU/ml), isothermal microcalorimetry and scanning electron microscopy. The colony counts after treatment with EDTA and DTT were similar to those after exposure to 0.9% saline (6.3, 6.1 and 6.0 log CFU/ml, respectively) for *S. epidermidis* biofilms, and (5.1, 5.2 and 5.0 log CFU/ml, respectively) for *P. aeruginosa* biofilm. Sonication detected higher CFU counts (7.5 log CFU/ml) for *S. epidermidis*; (p<0.05) and 6.5 log for *P. aeruginosa* biofilm (p <0.05). Concordant results were detected with isothermal microcalorimetry and scanning electron microscopy. In conclusion, the CFU count after treatment of *S. epidermidis* or *P. aeruginosa* biofilms with EDTA and DTT was similar to those observed after 0.9% saline used as control. In contrast, sonication was superior to chemical methods for biofilm dislodgment and detection of microorganisms in sonication fluid. In conclusion, our study showed that sonication is superior to chemical method to dislodge bacterial biofilm from artificial surface and should be considered as standard diagnostic method for biofilm detection in implant-associated infections.

## INTRODUCTION

Orthopedic devices are increasingly used for treatment of degenerative joint disease and for fixation of bone fractures. Infections represent a significant complication of implant surgery, resulting in major challenges in diagnosis and treatment. The crucial step in the management of orthopedic implant-associated infections is the accurate and timely diagnosis (1). However, this represents a considerable challenge, as these infections are caused by microorganisms embedded in a polymeric matrix attached to the device surface. In order to isolate and identify the causing microorganism, the dislodgment and dispersion of the sessile community represent the first step before plating the specimen on solid media (2). To improve biofilm removal from implant surface, different approaches had been described. Among others, sonication is based on a mechanical dislodgement (3), while treatments with the metal-chelating agent ethylenediaminetetraacetic acid (EDTA) (4) and the strong reducing agent, dithiothreitol (DTT) (5), might disgregate biofilm by chemical interactions.

Sonication of explanted components as an add-on procedure to routinely conducted microbiological analysis has shown to improve the pathogen detection (3), (6). Sonication reportedly yields rates of bacterial recovery from 70% to 100% compared to 10% to 100% when scraping the prosthetic surface and sensitivity of approximately 65% to 80% depending on prior antibiotic therapy (7), (8).

Recent reports considered the treatment of explanted prostheses with a solution containing DTT a potential alternative to sonication to dislodge biofilm-embedded bacteria and allow for subsequent isolation and identification of the microorganisms by conventional laboratory techniques (5). The ability of EDTA to chelate and potentiate the cell walls of bacteria and destabilize biofilms by sequestering calcium, magnesium, zinc, and iron suggests its use suitable for the biofilm detachment (4).

The aim of the study was to compare the ability of chemical (EDTA and DTT) and mechanical (sonication) methods alone or in combination to detach biofilm-embedded bacteria.

## MATERIALS AND METHODS

### Bacterial strains and biofilm growth conditions

Biofilms of *Staphylococcus epidermidis* (ATCC 35984) and *Pseudomonas aeruginosa* (ATCC 53278) were formed on porous glass beads (4 mm diameter, 60 μm pore sizes, ROBU^®^, Hattert, Germany). To form biofilms, beads were placed in 2 ml of brain heart infusion broth (BHIb, Sigma-Aldrich, St. Louis, MO, USA) containing 1×10^8^ CFU/ml bacterial inoculum and incubated at 37°C. After 24 h, beads were re-incubated in fresh BHIb and biofilms were let statically to grow for further 72 h at 37°C. After biofilm formation, beads were washed six times with 2 ml 0.9% saline to remove planktonic bacteria.

### Biofilm dislodgment by chemical methods (EDTA or DTT) or sonication

To define the minimal concentration and treatment duration for biofilm dislodging, washed beads were placed in 1 ml of EDTA at concentrations 12, 25 and 50 mM or DTT at concentrations 0.5, 1 and 5 g/l and exposed for 5, 15 and 30 min. Untreated beads incubated with 0.9% saline were used as negative control.

To evaluate the sonication effect, biofilms were sonicated as described previously (3). Briefly, each bead was inoculated in 1 ml 0.9% saline and sonicated at 40 kHz at intensity 0.1 Watt/cm^2^ (BactoSonic, BANDELIN electronic, Berlin, Germany) for 1 minute. One-hundred microliter of serial dilutions of the resulting sonication fluid or the solution obtained after chemical treatment with DTT or EDTA were plated onto Tryptic Soy Agar (TSA) (Sigma-Aldrich, St. Louis, MO, USA). After 24 h of incubation at 37°C, the CFU/ml number was counted.

Additionally, the viability of planktonic bacteria in presence of chemical agents was evaluated. Planktonic cells of *P. aeruginosa* and *S. epidermidis* at final concentration of ≈10^5^ CFU/ml were exposed to EDTA (25 mM) and DTT (1 g/l) for different time periods (5, 15 and 30 min). All experiments were performed in triplicates.

### Isothermal microcalorimetry analysis

To prove the dislodgment effect of previously described methods and reveal the presence of bacterial cells remained attached on the bead surface, treated beads were washed six times in 2 ml 0.9% saline to remove the dislodged biofilm and placed in 4 ml-glass ampules containing 3 ml of BHIb. The ampoules were air-tightly sealed and introduced into the microcalorimeter (TAM III, TA Instruments, Newcastle, DE, USA), first in the equilibration position for 15 min to reach 37°C and avoid heat disturbance in the measuring position. Heat flow (μW) was recorded up to 20 h. The calorimetric time to detection (TTD) was defined as the time from insertion of the ampoule into the calorimeter until the exponentially rising heat flow signal exceeded 100 μW to distinguish microbial heat production from the thermal background (9). Growth media without molds served as negative control.

### Scanning electron microscopy

Beads with biofilm were fixed with 2.5% (v/v) glutaraldehyde in natrium cacodylat buffer and the samples were dehydrated with increasing concentrations of ethanol for 2 min each. The samples were stored in vacuum until use. Prior to analysis by Field emission scanning electron microscope (DSM 982 GEMINI, Zeiss Oberkochen, Germany), the samples were subjected to gold sputtering (MED 020, BAL-TEC). All experiments were performed in triplicate.

### Statistic methods

Statistical analyses were performed using SigmaPlot (version 13.0; Systat Software, Chicago, IL, USA) and graphics using Prism (version 7.03; GraphPad, La Jolla, CA, USA). Quantitative data were presented as mean ± standard deviation (SD) or median and range, as appropriate. To compare different groups, nonparametric Kruskal-Wallis test and Wilcoxon signed-rank test for independent samples were used. The significance level in hypothesis testing was predetermined at p <0.05.

## RESULTS

### CFU counting method

The dislodged CFU counts after treatment of beads with *S. epidermidis* and *P. aeruginosa* biofilms at different concentrations and time points are shown in Figure 1 and 2. Evaluating dislodgement effect of chemical methods, mean colony count obtained after treatment of *S. epidermidis* biofilms with EDTA (25 mM, 15 min) and DTT (1 g/L, 15 min) was similar to those observed after 0.9% NaCl used as control (6.3, 6.1 and 6.0 log CFU/mL, respectively). By contrast, sonication detected significantly higher CFU counts with 7.5 log (p <0.05) (Figure 3A). Similar results were observed when *P. aeruginosa* biofilms were treated with chemicals (EDTA and DTT) or saline (5.1, 5.2 and 5.0 log CFU/ml, respectively). By using sonication, CFU count of 6.5 log (p <0.05) was observed. (Figure 3B).

**Fig. 1.**
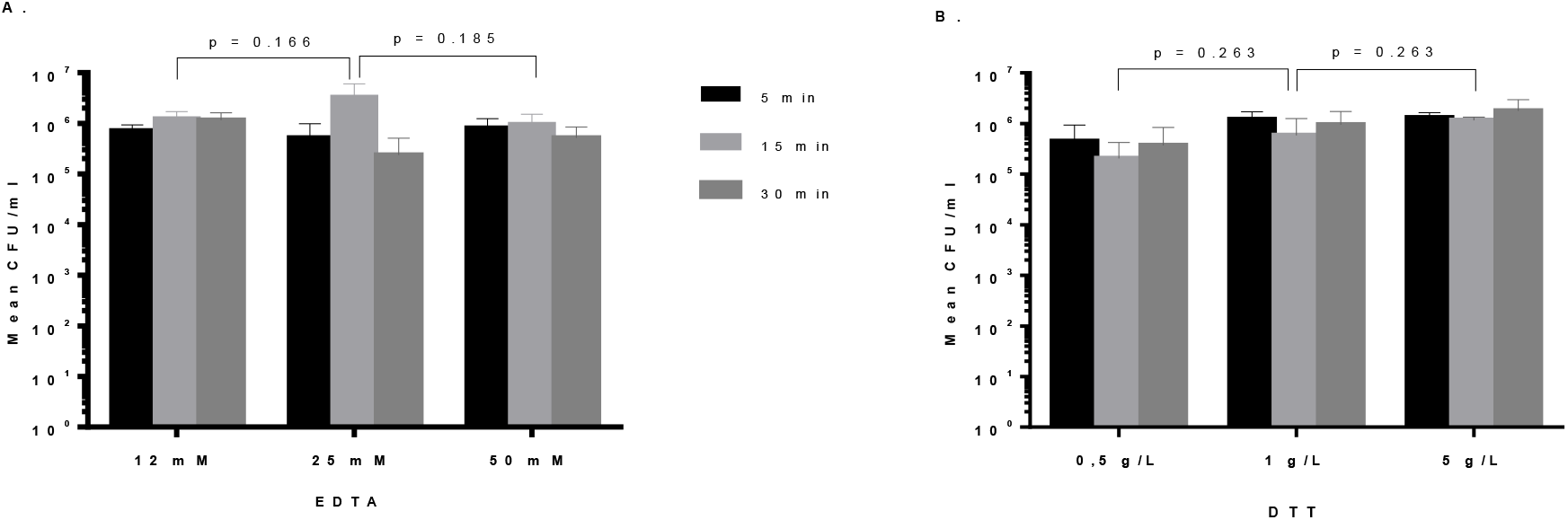
*S. epidermidis* biofilm after treatment with different concentrations of either EDTA (A) or DTT (B) at different time points. Mean values are shown, error bars represent standard deviation.

**Fig. 2.**
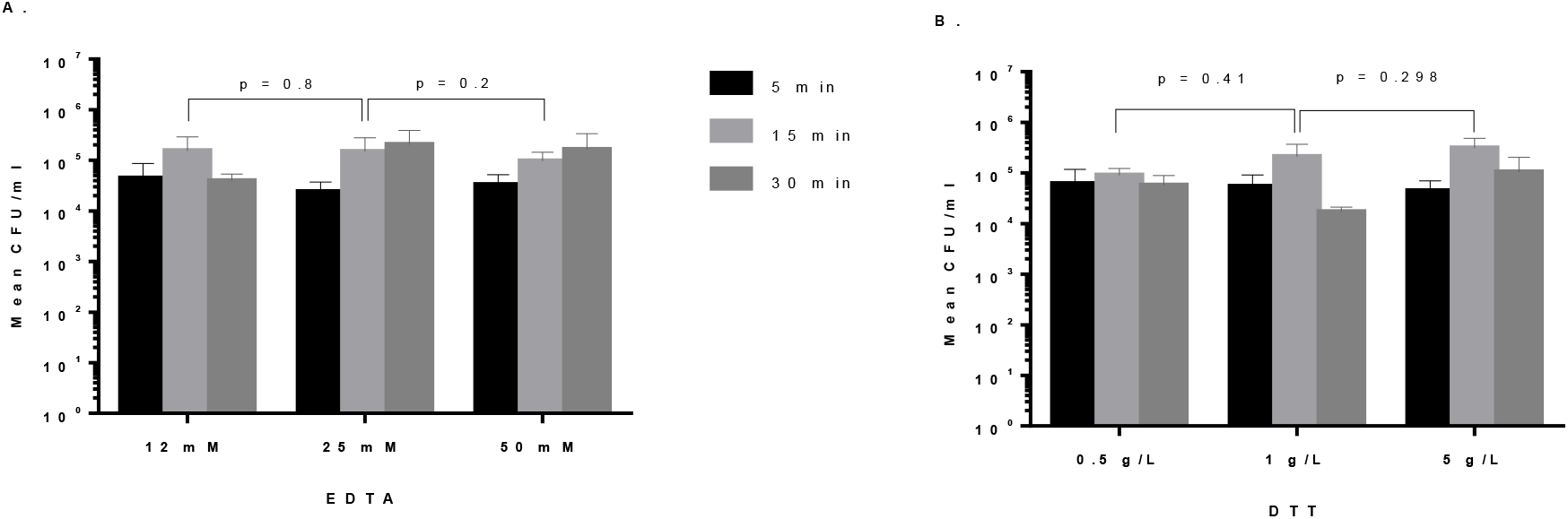
*P. aeruginosa* biofilm after treatment with different concentrations of either EDTA (A) or DTT (B) at different time points. Mean values are shown, error bars represent standard deviation.

**Fig. 3.**
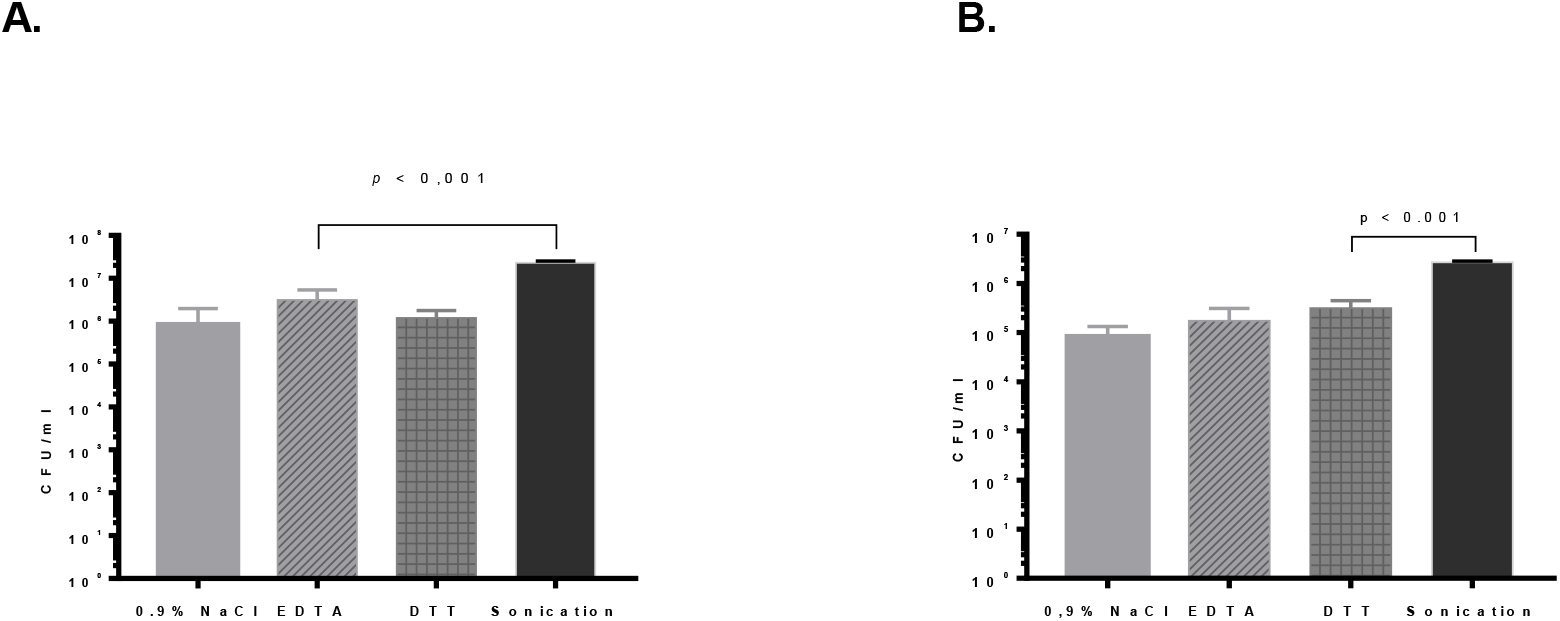
Quantitative analysis of biofilm dislodging methods. (A) *S. epidermidis* biofilm. (B) *P. aeruginosa* biofilm.

### Isothermal microcalorimetry

The heat produced by samples containing sonicated beads of *S. epidermidis* was detected after 11 h from monitoring start (100 μW was set as cut-off value). In contrast, heat production exceeding the threshold of 100 μW was observed earlier (after 6.3 and 6.5 h) for the samples that were previously treated with EDTA and DTT, confirming the presence of a higher number of residual bacteria on beads treated with chemical methods, in comparison to those after sonication. This difference was statistically significant (p <0.001). No difference in heat production was observed after treatment with 0.9% saline (control) and EDTA or DTT (6.3 vs 6.5 and 6.4 h, respectively) (p=0.3). Similar results were observed with the analysis of *P. aeruginosa* biofilm beads, although the time of heat detection after sonication of beads was significantly higher (11 h) in comparison to EDTA and DTT (6.5 and 6.5 h, respectively) (p <0.001); no difference between both chemical methods and the control (6.2 h) was observed (p=0.3) (Figure 4A and B).

**Fig. 4.**
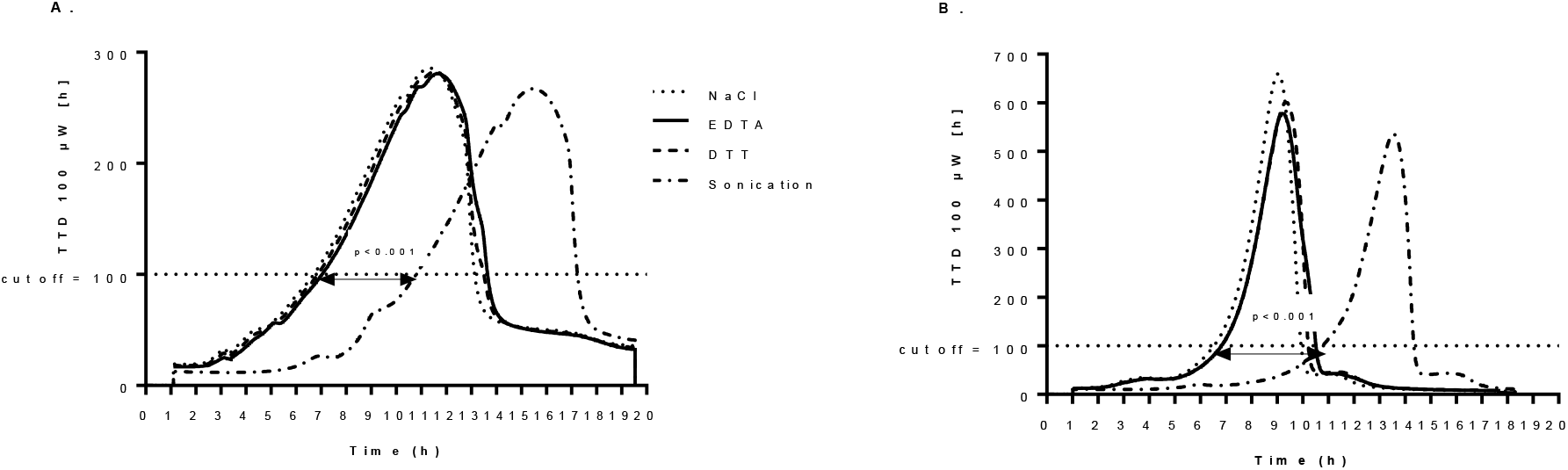
The microcalorimetric time to detection (TTD) a bacterial growth. A. *S. epidermidis* biofilm. B. *P. aeruginosa* biofilm.

### Scanning electron microscopy

The use of scanning electron microscopy (SEM) allowed to visualize the biofilms of *S. epidermidis* and *P. aeruginosa* before and after treatments with either chemicals or sonication. For both microorganisms the scanning electron microscope images showed substantial less biofilm biomass remaining on the beads when sonication was applied compared to control as well as both chemical methods (Figure 5 and 6).

**Fig. 5.**
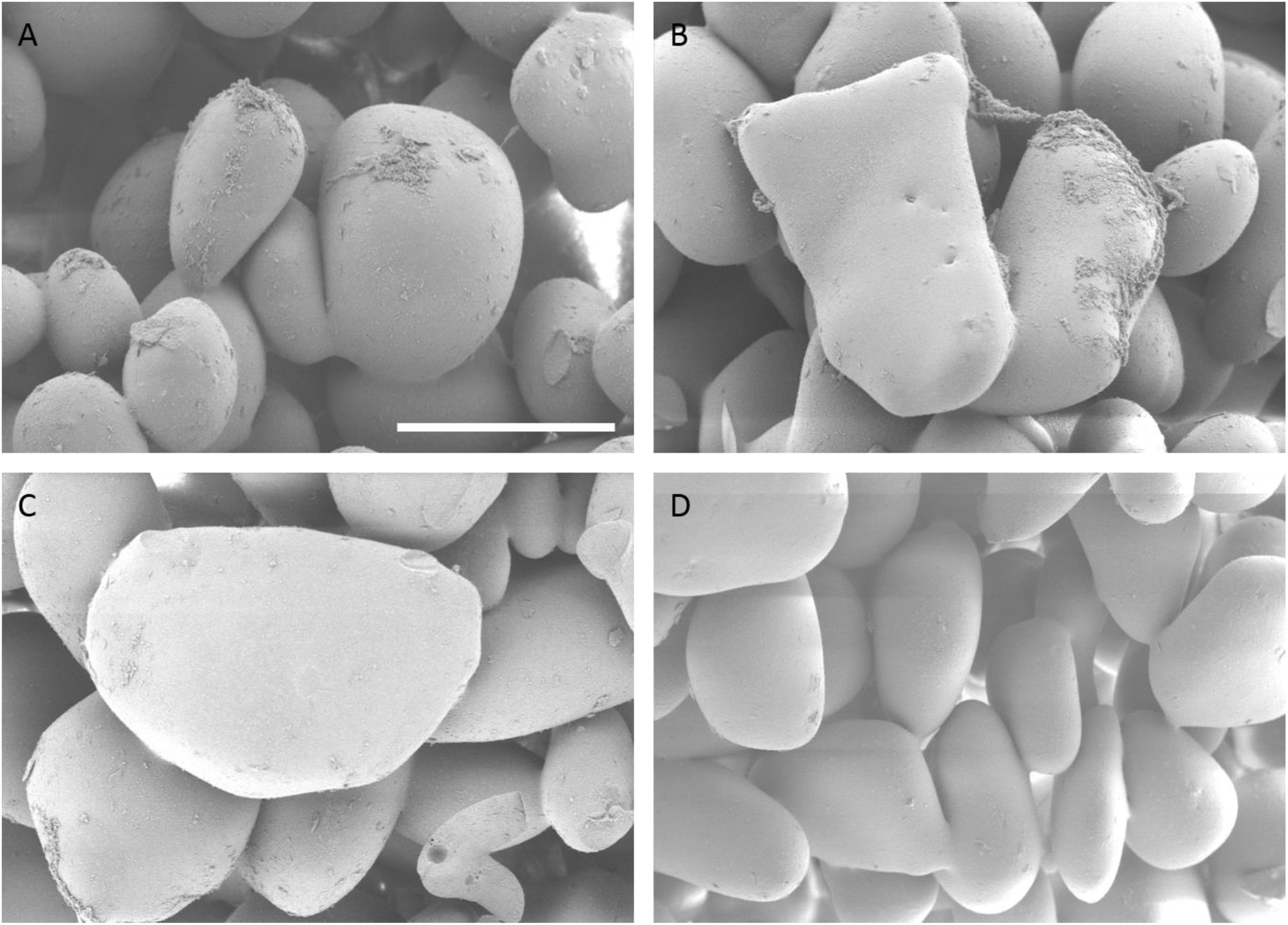
Scanning electron microscopy (SEM) of *S. epidermidis* biofilm: (A) beads after 0.9% saline treatment (control); (B) beads after EDTA treatment; (C) beads after DTT treatment; (D) beads after sonication treatment.

**Fig. 6.**
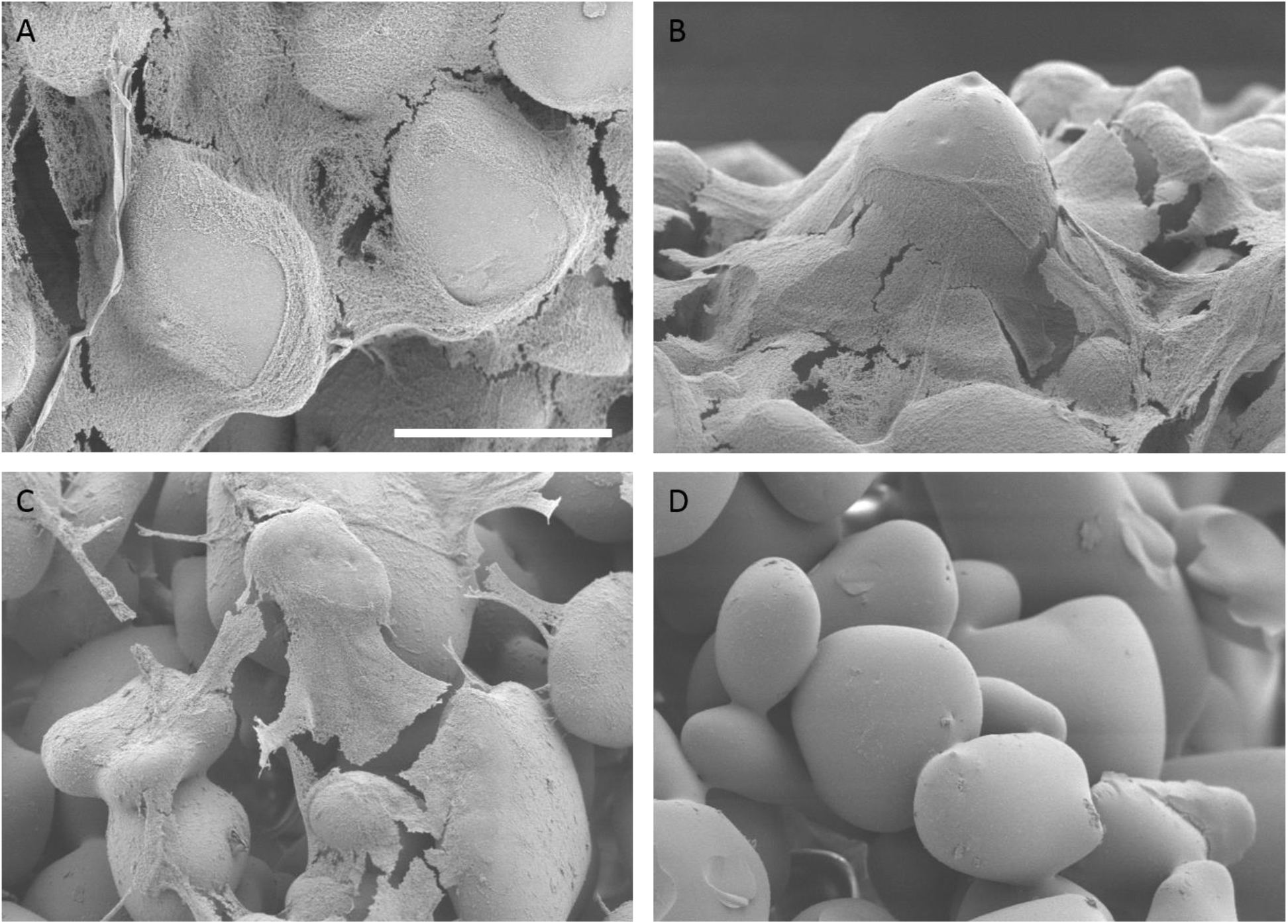
Scanning electron microscopy (SEM) of *P. aeruginosa* biofilm: (A) beads after 0.9% saline treatment (control); (B) beads after EDTA treatment; (C) beads after DTT treatment; (D) beads after sonication treatment.

## DISCUSSION

Implant-associated infections due to biofilm formation represent a major challenge for the microbiological diagnosis (10), (11). The presence of bacteria aggregated in a biofilm makes the detection of microorganisms challenging when the sample is seeded on standard medium, without any previous dislodgment and dispersion of the sessile community (2). We investigated the ability of different methods to dislodge *S. epidermidis* and *P. aeruginosa* biofilms from an abiotic surface *in vitro*, including sonication and chemical treatment with EDTA or DTT. To compare dislodgement effect of chemical methods, the concentrations of 25 mM EDTA and 1 g/l DTT were chosen as they showed significant increase in CFU count compared to other concentrations at the time point of 15 min. Concentration of 1 g/l DTT was also proposed by Drago et al. (5) The time of 15 min was chosen as a most appropriate time for the routine microbiological examination. (Fig. 1, Fig. 2).

Our results showed significantly higher dislodged CFU/ml for *S. epidermidis P. aeruginosa* biofilm when sonication method was applied in comparison to chemical agent DTT. No significant difference in the CFU number was observed after 1 mg/l DTT treatment, as compared to the control. Our results contradict those showed by other authors (5). In this study polyethylene and titanium discs covered with *P. aeruginosa, S. aureus, S. epidermidis* and *E. coli* biofilms were treated with DTT solutions at different concentrations and for different time points. The authors found that a solution of 1g/l DTT applied for 15 min was able to dislodge *P. aeruginosa* biofilm with similar yield as obtained by sonication, but the number of *S. aureus* biofilm cells removed by DTT were higher than that dislodged after sonication from the same materials. Similarly, colony numbers for S *epidermidis* were higher after DTT treatment than after sonication, whereas the number of *E. coli* colonies obtained after sonication and DTT were similar. The discordance in the study results might be explained by the influence of biomaterial type on the biofilm formation. In a recent study (12), the influence of biomaterials of retrieved hip and knee prosthesis on microbial detection by sonication was onstrateddem.

In our study EDTA was not able to efficiently dislodge bacterial biofilm from artificial surface. Cell colony count was similar to those obtained after treatment with 0.9% saline and significantly lower than those observed when sonication was applied. Previous reports demonstrated that EDTA affects *P. aeruginosa* biofilms (4), (13). Banin et al. suggested that exposure of *P. aeruginosa* biofilms to EDTA triggered detachment of cells from biofilms. They showed the dispersal of cells from biofilms caused by EDTA in a flow cell system. After addition of 50 mM EDTA to the medium reservoir, in 50 min they determined increase two log more in the number of cells in the effluent compared to untreated flow cell which showed a constant level of viable, dispersed cells in the effluent. They note that activity of EDTA in detachment of cells from the biofilm is mediated by chelation of several divalent cations such as magnesium, calcium, and iron that are required to stabilize the biofilm matrix.

Our results from counting the dislodged bacterial counts were confirmed by two additional independent techniques, namely isothermal microcalorimetry and SEM imaging. Isothermal microcalorimetry is a highly sensitive method for bacterial replication due to their metabolic heat production (14), (9). It has been widely employed in different studies to test the viability of either planktonic or biofilm bacteria after antibiotic treatment (15), (16). Here it was used to evaluate bacteria remaining on the glass beads after dislodging treatments. Isothermal microcalorimetry showed a significant delay in the detection of bacterial metabolism-related heat production from the beads with *S. epidermidis* when sonication was applied, as compared to chemical treatments - EDTA and DTT, suggesting that significantly less bacteria remained attached to the beads after sonication. Similar results were observed by the analysis of *P. aeruginosa* biofilm beads. The use of SEM allowed for visualization the biofilms of *S. epidermidis* and *P. aeruginosa* before and after treatments with either chemicals or sonication. In both types of bacterial biofilm SEM images showed less biofilm remaining on the beads when sonication was applied compared to the untreated control as well as both chemical methods.

In conclusion, our study showed that sonication is superior to chemical method for dislodgement of bacterial biofilm from surface and should be considered as the standard diagnostic method for biofilm detection in the diagnosis of implant-associated infection. Future studies may investigate a potential synergistic effect of sonication with chemical or mechanical dislodgement techniques, which do not affect the viability of microorganisms.

## ACKNOWLEDGMENTS

This work was supported by the PRO-IMPLANT Foundation, Berlin, Germany (https://www.pro-implant-foundation), a non-profit organization supporting research, education, global networking and care of patients with bone, joint or implant-associated infection. The funding had no influence on the data analysis or interpretation of the results.

